# Spatial Genome Organization as a Framework for Somatic Alterations in Human Cancer

**DOI:** 10.1101/179176

**Authors:** Kadir C. Akdemir, Yilong Li, Roel G. Verhaak, Rameen Beroukhim, Peter Cambell, Lynda Chin, P. Andrew Futreal, on behalf of the PCAWG-Structural Variation Working Group

## Abstract

Genomic material within the nucleus is folded into successive layers in order to package and organize the long string of linear DNA. This hierarchical level of folding is closely associated with transcriptional regulation and DNA replication. Microscopic studies concluded that genome organization inside the nucleus is not random and chromosomes within the nucleus create territories^1^. More recent chromosome conformation studies have revealed that mammalian chromosomes are structured into tissue-invariant topologically associating domains (TADs) where the DNA within a given domain interacts more frequently together than with regions in other domains^2,3^. Genes within the same TADs represent similar expression and histone-modification profiles^4^. In addition, enhancer-promoter pairs within the same TAD respond similarly to hormone induction^5^ or differentiation cues^6^. Therefore, regions separating different TADs (boundaries) have important roles in reinforcing the stability of these domain-wide features. Indeed, TAD boundary disruptions in human genetic disorders^7,8^ or human cancers lead to misregulation of certain genes^9,10^, due to *de novo* enhancer exposure to promoters. Here, to understand effects and distributions of somatic structural variations across TADs, we utilized single nucleotide variations, deletions, inversions, tandem-duplications and complex rearrangements from 2658 high-coverage whole genome sequencing data across various cancer types with paired normal samples. We comprehensively profiled structural variations with respect to their effect on TAD boundaries, on the regulation of genes in human cancers.

---

TADs are considered to represent functional domains because a given TAD encompasses the regulatory elements for the genes inside the same domain^11^. Therefore, integrity of the domain structures is important for proper gene regulation^12^. Boundary regions separating TADs are enriched for an array of proteins generally referred as architectural proteins^13^. Among the architectural proteins, insulator protein CCCTC-binding factor (CTCF) is the best-characterized the architectural protein^4^, along with the cohesin complex subunits. In addition, active transcription marks are enriched at TAD boundaries whereas heterochromatic histone marks such as H3K9me3 are depleted^4^. TAD boundaries are highly conserved among different cell types and different mammals at cell type-specific; yet still take place within encapsulating TAD structures^15^. Disruption of domain boundaries results in ectopic interactions between neighboring domains and impacts regulation of nearby genes^3,7^. Regulatory landscapes are an important part of human malignancies and studies have shown that enhancer “hijacking” can lead to overexpression of oncogenes (e.g. growth factor independent 1 family oncogenes [GFI1 and GFI1B]) in medulloblastoma^16^. Another example is the proto-oncogene *MECOM* (EVI1) that is exposed to regulatory elements within a different TAD as a result of an inversion event in acute myeloid leukemia (AML) cells. This alteration leads to an upregulation of the oncogene, which facilitates tumor formation^17^. Several other studies reported that likelihood of translocations between distal segments of DNA in cancer cells (such as IGH/MYC translocations in B-cell malignancies) is closely related with the three-dimensional spatial organization^18-20^. Hence, genomic rearrangements can play a significant role in reshuffling TAD structures, thereby resulting in altered gene regulation by the formation of aberrant long-range contacts between genes and regulatory elements.

## Characterization of a common set of TAD boundaries among different cell types

We utilized high-resolution chromosome conformation datasets (Hi-C) from five human cell lines representing three distinct embryonic germ layers (GM12878 and HMEC mesoderm; IMR90 endoderm; HUVEC and NHEK ectoderm)^21^ to identify TAD boundaries in different cell types (Supplementary Fig 1a). We called TAD boundaries from 25-kb resolution Hi-C data for each cell type with insulation score^22^ approach (Methods). This method calculates a score (TAD signal) for each 25kb bin’s average interactions with the nearby loci for a 2MB genomic window. Boundaries are determined as regions with local insulation minima along the diagonal of Hi-C matrix (Methods)^22^. As a result, a number of boundaries ranging from 3926 to 4690 in different cell types were found. We next asked whether our TAD boundary calls were consistent with the previously reported boundaries and showed attributes of TAD boundaries. To test this, we compared available boundary regions for IMR90 cells identified through a directionality-based approach (with 40Kb Hi-C resolution)^2^. Our IMR90 boundary calls were highly overlapping (> 84%) with published boundaries (Supplementary Fig 1b). This showed that current boundary regions were comparable with previously mapped boundaries even though they were identified with a different Hi-C resolution and detection algorithm. Furthermore, we observed known TAD-boundary signatures^2^ around our boundary calls for each cell type (Supplementary Fig 1c, Methods).

In order to use in this pan-cancer analysis, we sought to generate a common set of boundaries among the five profiled cell lines. There was a significant (p < 10^−6^) overlap between TAD boundaries among all profiled cell types and 2477 genomic sites were observed as common boundaries (Supplementary Table 1). Of note, the common boundaries set was the highest among all pair-wise comparisons (Supplementary Fig 1d). This observation confirmed previous studies that suggested a stable conformation of TADs^2-5,15^. The median distance between the common boundaries is approximately 750 kb, consistent with the reported median TAD size in human cells (Supplementary Fig 1e)^2,23^. The resulting 2477 common regions were used for the rest of the analyses (referred to as boundaries henceforth). Next, to test whether overall chromatin architecture is similar in cancer and non-cancer cells, we intersected these boundaries with the cancer cell line TAD boundaries. Using boundaries from a leukemia cell line K562 and a breast cancer cell line MCF7^24^ resulted in 85% and 83.4% overlaps, respectively (Supplementary Fig 1f-g). These analyses revealed a significant (p < 10^−7^) percentage of boundaries conserved between normal and malignant cells.

We examined enrichment of CTCF binding and DNaseI hypersensitivity (DHSs) sites, as well as active transcription start sites (TSSs) and heterochromatic regions around boundaries from various cell types previously profiled by the Encyclopedia of DNA elements (ENCODE) consortia^23^, and Roadmap Epigenome project^25^ (Methods). We observed that CTCF binding sites and active promoter marks were enriched, whereas heterochromatin state was depleted at the boundaries. In addition, TAD signal levels were lowest at the boundaries compared to flanking sites (Figure 1a), consistent with the role of TAD boundaries reducing the contacts between adjacent domains. Overall, these common 2477 boundaries exhibit the genomic features of TAD boundary across different human cell types.

**Figure 1.**
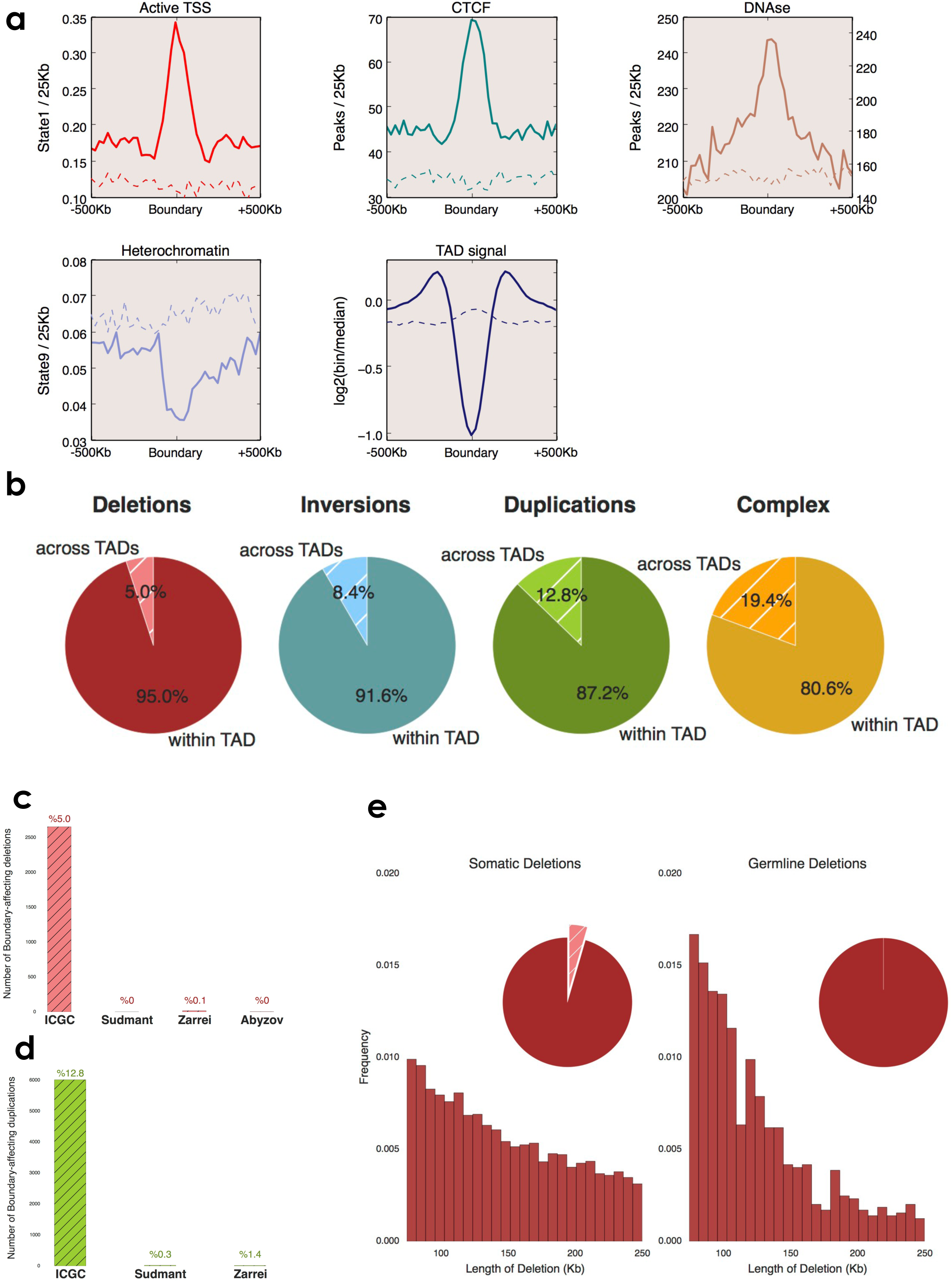
TAD boundaries are predominantly affected by somatic structural variations in cancer genomes. **a.** Profiles of active transcription start sites (TSS), CTCF peaks, DNaseI hypersensivity, heterochromatic states and TAD signal around common TAD boundaries. Dashed lines represent enrichment levels for the shuffled boundaries. **b.** Pie charts represent percentage of structural variants (length <= 2Mb) ‘across TADs’ (shaded) and ‘within TADs’ (solid) observed in cancer genomes. **c-d)** Bar charts show TAD-boundary affecting **c)** deletions and **d)** tandem-duplications in cancer genomes, and in genomes of healthy individuals from three different studies^25-27^. **e)** Histograms show the length distribution somatic and germline deletions in ICGC cohort with size range 75-250kb. Pie charts represent percentage of structural variants ‘across TADs’ (shaded) (%4.1 for somatic and less than %0.1 for germline deletions) and ‘within TADs’ (solid).

## TAD boundaries are frequently affected in human cancers

To understand the effects of structural variations (SVs) on TAD boundaries in human cancers, we used SV break-point orientations as a method to classify events as deletions, inversions, duplications or complex rearrangements. Complex rearrangements include chromotripsis^26^ and other alterations, which include concomitant regional deletions, inversions or duplications. Structural variations were further categorized into two subgroups based on the length of the events. SVs larger than 2MB of length (long-range SVs) and shorter than 2MB in genomic length (short-range SVs). A majority of deletions, inversions and duplications can be categorized as short-range, however, complex events tend to be larger in length (Supplementary Figure 1h) In this study, we primarily focused on short-range SVs because each long-range SV could affect at least one boundary due to the genomic length of the event. We identified SVs that impacted the TAD boundaries (SVIBs) as the ones that span through the whole length of a boundary (∼75Kb). As a result, 5.0%, 8.5%, 12.8% and 19.9% of all deletions, inversions, duplications and complex events are called as SVIBs, respectively (Figure 1b). These ratios are strongly enriched for duplications (p < 10^−4^) but not for deletions, inversions or complex events (p > 0.05) compared to the shuffled TAD boundaries (Methods). In cancer cells, boundaries are frequently affected due to frequent structural alterations; therefore, resultant SVIBs may play a role in carcinogenesis. Length distributions of the SVIBs are higher compared to all short-range SVs, yet still uniformly distributed (Supplementary Fig 1i). Most of the SVIBs target a single boundary, for all SVIB events 74% of deletions, 65% of inversions, 71% of duplications and 64% of complex events affect a single boundary per variations (Supplementary Fig 1j). SVIBs are distributed throughout the genome on all chromosomes and affect a majority (98.4%) of the boundaries, except a few boundaries located at the low mappability regions of the genome (Supplementary Fig 1g). TAD boundaries are significantly less likely (p < 0.02) to be affected by known deletion/duplication polymorphisms derived from healthy human populations^27-29^ (Figure 1c). Genomic length of the germline alterations tends to be shorter compared to somatic alterations observed in tumors due to negative selection against large SV in germline^30^. Therefore, we selected germline and somatic deletions between 75kb and 250kb genomic length occurring in ICGC samples (Figure 1d). This filtering ensured that selected somatic (median: 137Kb) or germline (median: 113Kb) deletions have the length potential to disrupt TAD boundaries. We observed that germline deletions affecting TAD boundaries are rare (less than 0.1%; 6 affecting out of total 924 deletions) compared to somatic deletions (4.1%) even with the similar genomic ranges, suggesting that germline variations in TAD boundaries may not be as well tolerated as similar to somatic alterations.

## Distribution of SVIB is cancer- and chromosome-specific

We next focused on the distributions of SVIBs across 38 different ICGC cancer types. Number of SVIBs is generally follow the total number of SVs in a given cancer types. Our analysis revealed that among all cancer types leiomyosarcoma and uterus adenocarcinoma having higher numbers, in average 25 and 22 SVIBs per sample respectively, compared to median ∼7 SVIBs per sample across all ICGC cohort (Figure 2a-b). Ovarian, esophagus and breast adenocarcinoma cancers also contain high numbers of SVIBs, in average 20, 19 and 18 SVIBs per sample respectively. On the other hand, hematopoietic cancers (myeloid-MDS or myeloid-AML) demonstrated the lowest SVIB rates. Only glioblastoma samples (CNS-GBM) showed less than expected SVIBs (p<10^−3^) across all cancer types. Median SVs length in a given cancer type is not strongly correlated with the observed distributions (r^2^: 0.03-0.45) (Supplementary Fig 2a). Long-range SVIBs show similar distributions across cancer types where leiomyosarcoma and breast adenocarcinoma again containing higher number of SVIBs compared to other cancer types and leukemia samples having no SVIBs per sample (Supplementary Fig 2b). Taken together, our findings demonstrated that the impact of SVIBs is substantially varied across tumor types and thus may play a role in cancer type specific manner. Next, we sought to identify recurrently affected boundaries in specific cancer types. Of those recurrently affected boundaries (Supplementary Table 2, Methods), two adjacent boundaries between *KIAA1549* and *BRAF* genes are prone to tandem duplications specifically in pilocytic astrocytoma samples (Figure 2c). This region has previously been implicated in pilocytic astrocytoma, producing an oncogenic fusion between the aforementioned genes^31^. A boundary partially overlapping with an isoform of an alternative splicing regulator (*RBFOX1)*^32^ on chromosome 16 is deleted in 31 % of colorectal adenocarcinoma samples suggesting a potential function of this deletion in colorectal cancers (Figure 2d). A boundary on chromosome 3 overlapping with TUSC7 gene is recurrently deleted in osteosarcoma (15 %) but also in ovarian adenocarcinoma (12 %) (Supplementary Fig 2c). In addition to short-range SVIBs, two boundaries between *ERG* and *TMPRSS2* genes are most recurrently deleted by long-range SVs in prostate adenocarcinoma samples (Supplementary Fig 2d), agreeing with the known gene-fusion event between these two genes^33^. We surveyed the SVIB distributions on individual chromosomes and observed that SVIBs correlated with the number of boundaries on a given chromosome (r^2^: 0.68 – 0.92) (Supplementary Fig 2e-f). As well, gene density correlates with the number of SVIBs (r^2^: 0.7 - 0.85). Notably, distributions of SVIBs per chromosome are generally cancer specific, such as chromosome 12 is most affected in osteosarcoma likely due to neochromosome formations including MDM2 and CDK4 genes^34^, whereas chromosome 17 is affected in breast and esophageal adenocarcinoma samples predominantly by complex events (Supplementary Fig 2g). In addition to higher level of SVIBs, we observed a higher mutational load specifically on chromosome 12 in osteosarcoma samples (Supplementary Fig 2h). These findings emphasize the cancer-specificity of SVIBs, where active mechanisms leading to overall SV burden and structure in different tumor types yield potential changes in TAD structures in each cancer type. We examined SVs occur within TADs which result in disruption of CTCF-based “insulated neighborhoods”^35^. We identified a set of insulated neighborhood disruptions in various cancer types, such as recurrent loops perturbations in esophageal, gastric and colon adenocarcinoma involve a neighborhood encapsulating the *FOXC1* (Figure 2e), or another neighborhood near to the *CLCN4* where a CTCF site is deleted (Figure 2f). Other recurrently altered neighborhoods include a CTCF site near to the *BCL6* in hepatocellular carcinoma and breast adenocarcinoma (Supplementary Fig 2i, Supplementary Table 3). Chromatin folding perturbations are not specific to TAD boundaries and are observed at the level of CTCF-CTCF chromatin loops, therefore chromatin misfoldings occur at various scales in cancer genomes.

**Figure 2.**
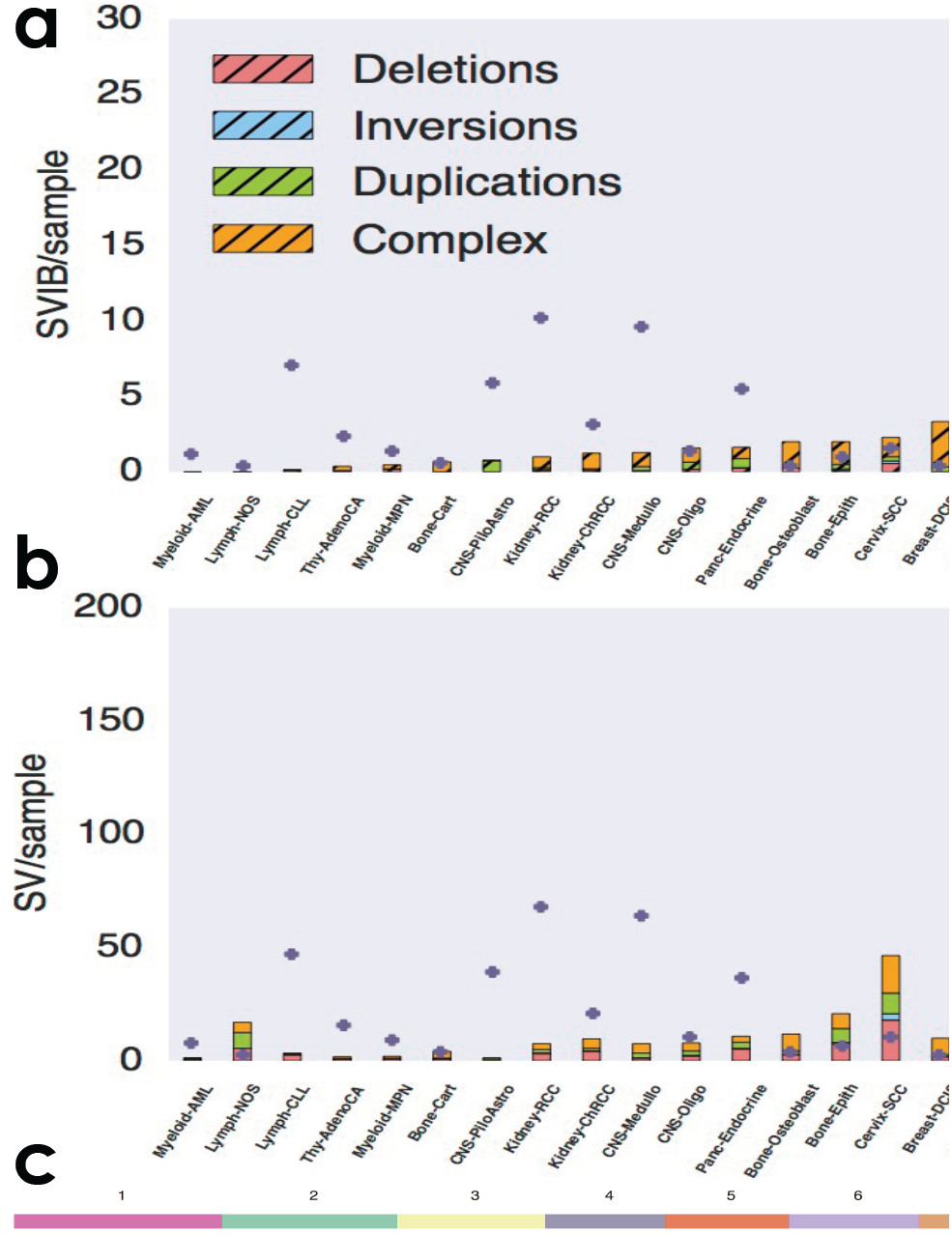
Genome folding disruptions are cancer specific. a-b) **a.** Bar charts represent the distribution of average number of SVs (length <= 2Mb) that impacted the TAD boundaries (SVIBs) per sample for each histology type. **b.** Bar charts show the distribution of average number of SVs (length <= 2Mb) observed in each histology type. Yellow dots represent patient numbers for each study. Deletion, inversions, tandem-duplications and complex rearrangements are represented with red, cyan, green and orange colors, respectively. **c.** Recurrently duplicated adjacent TAD boundaries in pilocytic astrocytoma. Colored bars on top depict chromosomal locations of the boundaries. Columns of the heatmap are TAD boundaries and rows represent each pilocytic astrocytoma sample. TAD boundary affecting tandem-duplications are colored in green. Schematic below shows duplicated two adjacent boundaries (green boxes) between KIAA1549 and BRAF genes. **d.** A recurrently deleted TAD boundary in colon adenocarcinoma samples. Colored bars on top depict chromosomal locations of the boundaries. Columns of the heatmap are TAD boundaries and rows represent each colon adenocarcinoma sample. TAD boundary affecting deletions are colored in red. Schematic below shows deleted boundary (red box) partially overlapping with *RBFOX1*. **e-f.** Recurrently deleted insulated neighborhoods in esophageal, gastric and colon adenocarcinoma (**e.)** a CTCF-binding site near to *FOXC1* and (**f.**) a CTCF-binding site near to *CLCN4*. Black boxes show TAD boundaries, arcs represent CTCF ChIA-PET loops (gray). CTCF ChIP-Seq signal is represented by purple histogram. Vertical bars depict deletions in individual samples from esophageal, gastric and colon adenocarcinoma.

## Deletions between different epigenetically-enriched TADs can result in aberrant gene-expression changes

To ascribe potential functional effects of SVIBs on chromatin domains, we annotated the TADs by profiling the context of aggregate chromatin states within each TAD. We utilized a probabilistic approach calculating the occurrence of chromatin states in Roadmap Epigenome cell types. Coverage of 15 chromatin state enrichments in each domain was calculated and normalized to the length of the domain (Methods). The obtained matrix was grouped with k-means clustering approach and five distinct groups of TADs were identified similar to the recent classification of chromatin domains^21-23,36^. These groups comprise heterochromatin (61), low/quiescent (705), repressed (481), low-active (764) and active (365) domains (Figure 3a, Supplementary Table 4). Next, we evaluated the annotation results by profiling the distributions of domain sizes. Repressed domains were larger in size and covered majority of the genome compared to active domains, in agreement with earlier TAD annotations (Supplementary Fig 3a-b)^23,37^. Median expression of the genes within each domain was calculated for 2921 cancer-free samples from 45 different tissues (GTEx consortium)^38^ as well as 998 cancer patients from ICGC expression datasets. Analysis of expression levels confirmed that genes within repressed domains had a significantly lower expression patterns than genes within active domains (p < 2.2^-16^) (Figure 3b, Supplementary Fig 3c). Finally, distributions of replication timing from various cell types and open/closed chromatin compartment calls from TCGA data^39^ corroborated with the annotated domains (Supplementary Fig 3d-e, Methods). Utilizing our domain annotations, we checked the distributions of flanking domains for deletion, inversion or duplication SVIBs. The majority of the SVIBs affected same flanking domain types such as boundaries separating low to low domains or low-active to low-active domains (Supplementary Fig 3f). Low to low-active domain boundaries were among the most highly affected boundaries, presumably because these two boundary types tend to be neighbors within the genome (Supplementary Fig 3f). Gene expression values were available for a subset of cancer samples. Therefore, we compared expression values of the genes that are residing on each side of the structural variations. We specifically focused on deletions affecting repressed and active domains as recent studies showed that fused repressed-active domains could lead to an up-regulation of nearby genes^40,41^. Indeed, genes located on the repressed side of deletions were significantly up regulated (p < 0.001, Wilcoxon-rank, Supplementary Table 5) in the sample harboring deletion compared to the rest of the samples in the same cancer study (Figure 3c). On average, repressed domain genes are up-regulated 5.02-fold (fold changes range between 71-to 0.8-fold) but active domain genes expression change 1.42-fold (fold changes range between 5-to 0.2-fold) compared to unaffected samples (Supplementary Table 5). However, the same effect was not observed for the genes residing on the active domain side of the deletion (Supplementary Fig 3g). This observation confirms the importance of TAD boundaries for reinforcing the separation of features between the different domains types. For example, a deletion in a malignant lymphoma sample was associated with a 37-fold increase *WNT4* level compared to the rest of the lymphoma patients. However, no expression change was observed for the active domain gene, *RPL11* (Figure 3d). Similarly, a deletion in a breast adenocarcinoma patient genome correlated with a 26-fold over-expression of *SLC22A2* compared to the rest of the breast cancer patients (Figure 3e). These results suggest that the effect of ectopic interactions upon SVIB deletions would generally be reflected on the repressed domain genes compared to the active domain genes.

**Figure 3.**
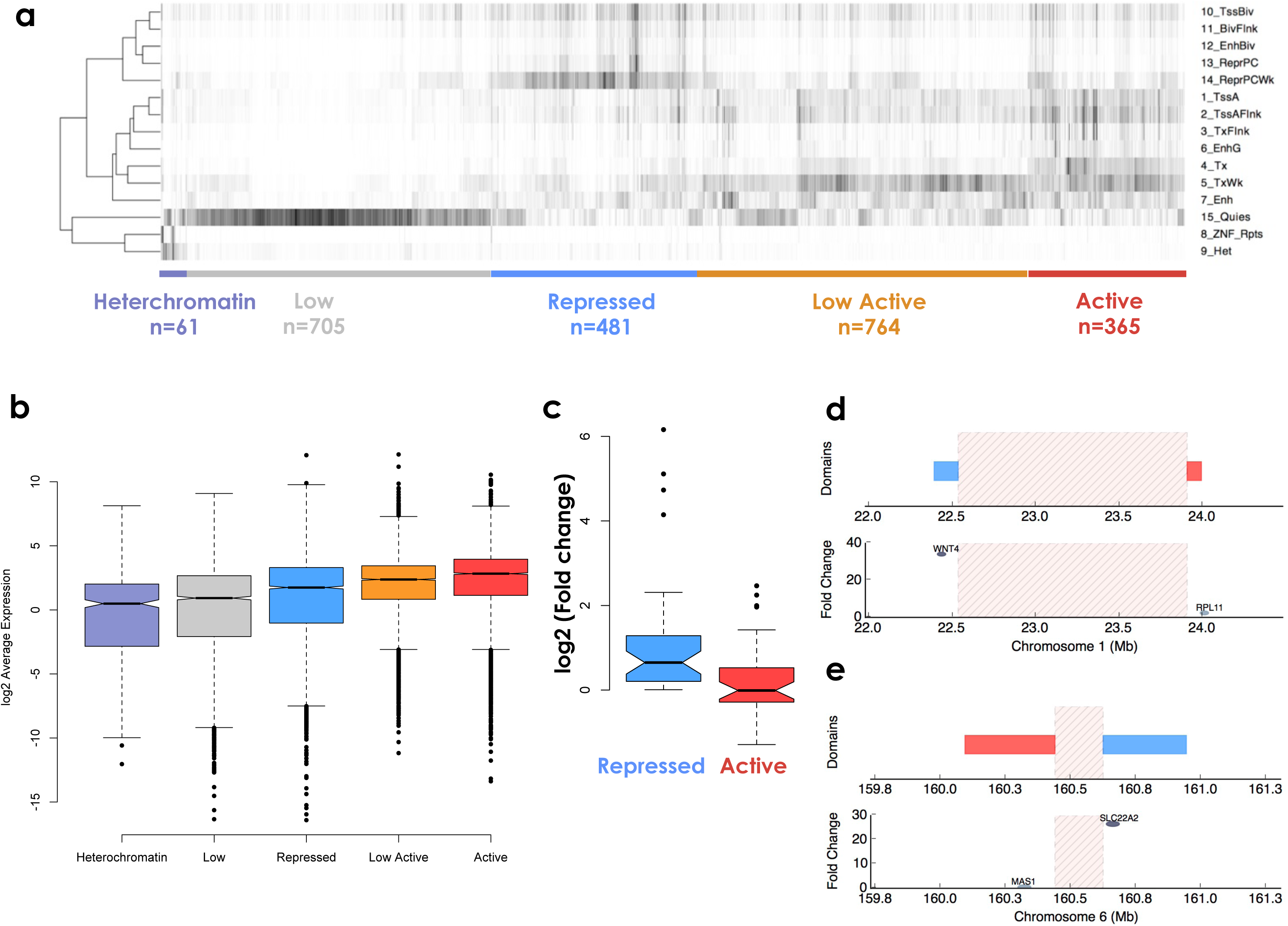
Deletions between active and repressed domains results in up-regulation of the repressed-domain genes. **a.** Classification of TADs based on chromatin state coverage. Heatmap shows domain-length normalized coverage of each chromatin state from Roadmap Epigenome aggregate data (rows) for each domain (columns). Chromatin states are clustered based on co-enrichments within TADs. Domains are classified into five groups: heterochromatin (purple), low/quiescent (gray), repressed (blue), low-active (orange), and active (red) according to dominant chromatin state combinations. Numbers show total number of TADs identified for each group. **b.** Box plots represent log2 average expression levels for genes located in each domain annotations. Average expression levels were calculated across all ICGC samples. **c.** Box plots show log2 fold-change for genes nearest to deletion break-ends on repressed TAD (blue) or active TAD (red) side of the deletions. Fold change was calculated based on the gene’s expression in the sample harboring the deletion compared to the rest of the samples in the same cancer study. **d.** An example of repressed gene overexpression upon boundary deletion in a lymphoma sample. Highlighted area represents the deleted region in the lymphoma sample. Tiles show flanking repressed (blue) and active (red) TADs to the deletion. Dots show fold-change in gene expression for *WNT4* (>37 fold) and *RPL11* genes in the boundary-affected lymphoma sample. **e.** An example of overexpression of a repressed gene upon boundary deletion in a breast cancer sample. Highlighted area represents the deleted region in the breast cancer sample. Tiles show flanking repressed (blue) and active (red) TADs to the deletion. Dots show fold-change in gene expression for *SLC22A2* (>26 fold) and *MAS1* genes in the boundary-affected breast cancer sample.

## Mutation load in human cancers is correlated with TADs

Finally, to elucidate if there is a connection between regional mutation rates and spatial genome organization, we profiled the distribution of somatic mutation rates from ∼51.5 million mutation calls across 2738 ICGC cancer samples with respect to TAD annotations (Methods). A higher mutational load was observed in heterochromatic regions, whereas active chromatin regions harbor fewer mutations as reported earlier^42-44^. Next, we used average replication timing data (Repli-Seq signals) from different ENCODE consortia cell types as a proxy for approximate replication timing in human cells. The number of mutations and the average replication timing of a given domain showed a negative correlation for each domain category (r^2^ between -0.48 to -0.68) (Figure 4a, Supplementary Fig 4a). Besides, mutation load and replication timing differed drastically across the TAD boundaries if neighboring TADs were epigenetically distinct, e.g., boundaries that demarcate low-active to low domains (black-arrow) (Figure 4b, Supplementary Fig 4b). However, mutational load did not change if a boundary was separating the same type of domains, such as low-active to low-active (orange-arrow) domains (Figure 4b). A recent study demonstrated that replication domain boundaries align with TAD boundaries therefore replication domains are regulated as the unit of TADs^45^. Here, we postulate that mutational load in cancer genomes is coinciding with TAD units as well, which could explain differences in regional mutation loads flanking the TAD boundaries. This idea prompted us to investigate the average distribution of mutations on either side of TAD boundaries between epigenetically distinct and similar TADs. Indeed, average mutation load substantially increased downstream the boundaries from low-active to low domains, while the replication signal declines (Figure 4c). However, domains separating epigenetically similar TADs (low to low or low-active to low-active) did not show any significant change around the boundaries for either mutational load or replication timing (Figure 4c). A similar trend of decrease in mutation load was also observed at the boundaries separating repressed to active or repressed to low-active TADs, indicating that mutation load is higher at epigenetically inactive TADs compared to the more active TADs (Supplementary Fig 4c-e). These results highlight the implicit link between spatial organization of chromatin and regional mutation loads in cancer cells.

**Figure 4.**
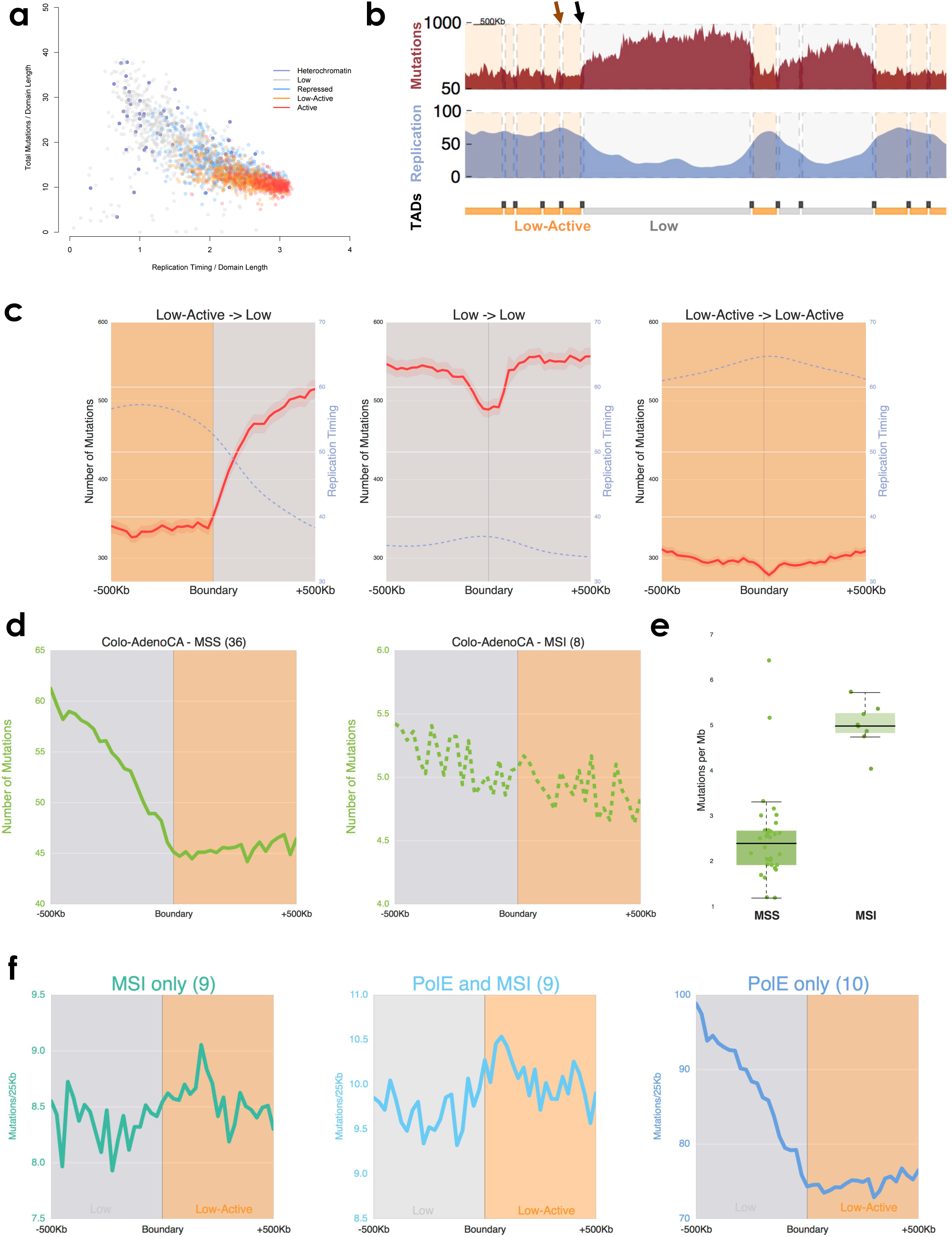
Somatic mutation distributions are correlated with the spatial chromatin organization. **a.** Scatter plot shows mutation loads (y-axis) and average replication timing (x-axis) for each domain. Colors represent distinct TAD types. **b.** An example region shows the correlation between mutation distributions and TAD types. Mutation load (red) and replication timing (blue) across different TADs are plotted on chromosome 11 between 75-84Mb. Tiles (below) show domain types (low TAD: gray, low-active TAD: orange) and black bars denote TAD boundaries. Shaded area represents designated domain types for genomic regions. **c.** Mutational load is correlated with spatial chromatin organization. Average profiles of mutation load in all ICGC samples (red) and replication-timing (purple) across 500Kb of boundaries delineating: low-active (orange) to low (gray); low (gray) to low (gray); and low-active (orange) to low-active (orange) TADs. **d.** Mutational load is flatter in tumors with miscrosatellite instability (MSI) across the TAD boundaries. Aggregate plots show sum of mutation loads between low TADs (gray) and low-active TADs (orange) for MSS-colorectal samples (Colo-AdenoCA-mss) (solid line) and MSI-colorectal samples (Colo-AdenoCA-MSI) (dashed line). Sample numbers are denoted on top of each plot. **e.** Boxplot demonstrates the mutations per megabase in each MSS (excluding PolE-deficiency samples) and MSI colon adenocarcinoma samples. **f.** Average profiles of mutation loads for only MSI (left), MSI and PolE-deficient (middle) and only PolE-deficient (right) samples across boundaries delineating low to low-active domains. Sample numbers are denoted on top of each plot.

Given that heterochromatic domains are generally positioned near the nuclear periphery, constitute condensed structures and late replicating, observed mutation load increase at these TADs were expected to be a result of inefficient DNA repair^46,47^. To examine whether the observed switch at the boundaries was diminished in cancers with known DNA repair deficiencies, we analyzed colon adenocarcinoma cases as DNA-repair pathway (MMR) deficiencies frequently observed in this tumor type. As a result, 8 samples were identified where mutation profiles were much less changed across TAD boundaries and exhibited nearly flat profiles compared to remaining 37 samples (Figure 4d). We annotated these 8 samples as mismatch repair deficient (MSI-high) samples compared to 37 mismatch repair proficient (MSS) cases. To assess our MSI calls, we utilized the results of a study that focused on mutational load with respect to replication timing^42^. This study categorized all of our flat-profile samples as MSI phenotype^42^, conforming our current findings by an independent approach. To more systematically identify MSI cases within the ICGC cohort, we classified all 2738 samples by comparing mutation loads across boundaries separating low domains from low-active domains (Methods, Supplementary Fig 4f). An additional 12 cancer samples were identified as MSI, from uterus (4), stomach (5), pancreas (1), liver (1) and glioblastoma (1) cancer types (Supplementary Fig 4g). These MSI identified samples indeed exhibit enrichment of mutational signatures attributed to DNA mismatch repair deficiency (Supplementary Figure 4h). Furthermore, we profiled the mutation distribution in primary and recurrent cancers with hereditary biallelic MSI deficiency syndrome^48^ and observed the similar flatter distributions across the boundaries. Therefore, both acquired and inherited MSI lead to similar flatter mutational distributions in cancer genomes (Supplementary Fig 4i, Methods). Overall, MSI samples harbor a higher somatic mutation burden compared to MSS samples (Figure 4e). Hence, we investigated samples with mutations in the proofreading domain of DNA polymerase ε (Polε) because this kind of DNA repair deficiency was observed concurrently with MSI in human cancers^49^ and Polε-deficiency generates high somatic mutational load in cancer genomes. Samples with Polε mutations exhibited a canonical mutational distribution observed in MSS samples despite having a higher mutational load compared to MSI samples (Figure 4f). Thus, a high level of mutation load is not the reason for the observed MSI-type mutation distributions. However, the samples categorized as both MSI and Polε-mutated^49^ showed an MSI-type flatter mutational distribution pattern, suggesting MSI events are the cause for the observed flatter mutational distributions (Figure 4f). Thus, regional mutation variation along TADs in cancer genomes is correlated with TAD types and proficiency of DNA repair complexes and there may well be a hierarchy of repair effects that are more or less prone to be influenced by TADs.

## Discussion

In this study, we explored the distributions of somatic genomic variations in human cancers with respect to TADs. We found that certain boundaries are affected in a cancer-specific manner and that chromosomes harboring higher number of genes are more prone to SVs and thus to more TAD-affecting SVs. In addition, we observed that structural variations in cancer cells predominantly affect TAD boundaries, which resulted in aberrant interactions between flanking domains, and potentially contributes to re-shaping gene expression and mutation profiles around the inflicted regions. Gene expression patterns are susceptible to changes due to *de novo* enhancer exposure of repressed genes in a variety of cancer types, therefore observed changes are likely due to chromatin domain reconfiguration instead of a cancer specific mechanism. The availability of tissue-specific high-resolution chromatin organization datasets and chromatin states will augment our understanding of structural variations and their impact on genome folding and transcriptional dysregulation in cancers^50,51^.

Furthermore, we report here that regional mutation distribution of human cancers is affected by spatial organization of the chromatin. Mutation loads changed across TAD boundaries, which suggest a link between genome folding, efficiency of DNA repair mechanisms and mutagen exposure. We found that DNA mismatch repair deficient-cancers did not display a similar mutational distribution. We utilized this flat signature of mutations across low and low-active TAD-separating boundaries and successfully identified several cancer samples that displayed an MSI phenotype. Hence, this approach can be used to detect MSI cases without any prior information about the genetic and epigenetic status of factors that have been functionally linked with the MSI occurrence. Architectural proteins are an important part of hierarchical chromatin organization, we checked whether CTCF-, RAD21- or SMC3-mutated samples show aberrantly high structural variations or different mutational distributions across TADs but we did not observe any drastic changes compared to the rest of the samples. Taken together, our study revealed a strong link between spatial organization of the genome inside the nucleus and oncogenic events such as structural variations and somatic mutations in human cancers.

## Methods

### Hi-C data analysis

Chromatin conformation assay (Hi-C) data for cell lines of GM12878, HUVEC, IMR90, HMEC, NHEK and K562 were downloaded from GEO (GSE63525). Intra-chromosomal 25 Kilobase-resolution raw observed, MAPQGE30 filtered values were normalized by diving with the multiplication of Knight and Ruiz normalization scores for two contacting loci. We calculated TAD signal by moving a window across the Hi-C matrix diagonal and calculated sum of interaction for a given bin up to 2 Mb flanking regions and calculated log2 of observed bin to mean of interaction values within the given 2 Mb window. To identify TAD boundaries, we utilized an approached which is based on insulation score calculation^22^, and called TAD boundaries for each chromosomes of each cell line with the following parameters: “-is 1000000 -ids 200000 -im mean -bmoe 1 -nt 0.1 –v”.

To calculate the significance of overlap between different TAD boundary calls, we converted the boundary regions into binary bins per genome to compare the overlap between previously published IMR90 TAD boundaries^2^ with our IMR90 boundary calls. We performed logical AND operation where if only two bins for the same genomic location of each condition are 1 then the region is counted as overlapping boundaries between two datasets. We used bootstrapping to determine distribution of the random overlap numbers between two calls, and calculated p-values based on observed number and distribution from shuffled boundaries. Shuffled boundaries are generated by randomly assigning boundaries with keeping the number of boundaries per chromosome constant. Obtained shuffled boundaries also converted to binary string and same logical AND operation applied. Shuffling performed for 10,000 times for a give boundary set. This procedure is applied in the rest of our study to generate shuffled boundaries. Next, we computed cumulative distribution of expected overlaps, z-scores and p-values were calculated based on observed number and obtained distribution from bootstrapping.

Common TAD boundaries are identified where boundaries of all five cell-types (GM12878, HUVEC, IMR90, HMEC and NHEK) are occurred within two Hi-C bins or 50 Kilobase in genomic range. Same bootstrapping method (explained above) was applied to calculate the significance of the overlap between common boundaries with TAD boundaries from cancer cell lines K562 and MCF7.

In order to cluster individual TADs (defined as genomic regions between two adjacent common boundaries) based on epigenetic modifications, we sought to utilize a comprehensive epigenome profiling from various human cell types. To this end, we used an entropy based approach (epilogos) to calculate occurrence of each chromatin state enrichment for a given genomic regions across all Roadmap Epigenome Consortia profiled cell types (http://compbio.mit.edu/epilogos/). We calculated the ratio of a TAD genomic space covered by each chromatin state, divided by the length of the TAD, and generated a normalized matrix where columns are TADs and rows are each chromatin states extensively studied by the Roadmap Epigenome Consortia^25^. We applied hierarchical clustering to rows to identify similar chromatin states and k-means clustering to columns to group TADs containing similar epigenetic modifications. We performed k-means clustering with k=2-8 clusters and decided on k=5 clusters as previous chromatin studies^21,23^ reveal 5 distinct epigenetically modified chromosomal domains and k=5 corresponded to better visually discernible domains. To determine how our TAD clustering correlate with gene expression in cancer-free and cancerous tissues, we downloaded normalized gene expression values for 2663 different cancer-free samples from GTEx Portal^52^ (v.1.6) and utilized normalized gene expression values for ICGC cancer samples. We plotted median expression of the genes in GTEx and ICGC samples, located in each domain type. Expression differences between heterochromatin and repressed domain expression with active domain expression were tested with Mann-Whitney test. We also calculated the total number of genomic regions covered by each domain type and plotted the ratios in Figure 3b and Supplementary Fig 3C. Finally, a recent report^39^ identified open and closed chromatin compartments (in 100Kb resolution) in cancer samples by using DNA methylation levels. We determined the percent of our domain calls covered with the open and closed chromatin calls from available cancer types.

We used HiCPlotter^53^ to plot Hi-C data with different features, TAD boundaries (Supplementary Fig1a) or gene-expression fold-changes after deletion between repressed and active domains (Figure 3c-d).

### Encode and Roadmap data

ENCODE replication timing data were downloaded from the UCSC Genome Browser ENCODE portal for the following cell types: BJ, GM06990, GM12801, GM12812, GM12813, GM12878, HeLa-S3, HepG2, HUVEC, IMR-90, K-562, MCF-7, NHEK and SK-N-SH. Replication timing values for smoothed wavelength transformed data were binned into 25Kb windows across the genome to discretize the data. Average of each bin values across all cell types were calculated and used as average replication timing in throughout the study.

We downloaded CTCF binding sites and DNaseI hypersensitivity for five cell types (GM12878, HUVEC, IMR90, HMEC, NHEK) from UCSC genome browser Encode portal. In addition, H3K9me3 and Input DNA ChIP-Seq alignment files (.bam) for each cell type were also downloaded. We randomly selected same number of alignment reads for H3K9me3 and Input DNA from .bam files and calculated log2 enrichment levels of H3K9me3 over Input DNA.

We downloaded all available CTCF peak calling results and DNaseI hypersensitivity regions from UCSC Genome Browser ENCODE portal from 80 and 115 different cell lines respectively (Supplementary Table). Occurrences of CTCF and DHSs per 25Kb windows across the genome are calculated for all downloaded cell types and used to calculate for TAD boundary and shuffled boundary enrichments.

### Structural Alterations

We downloaded ICGC consensus structural variations and annotations of each alteration (deletions, inversion, duplications, complex rearrangements) results (Li Y et al., Patterns of structural variation in human cancer). We specifically focused on the events occurring within a chromosome in this study, thereby did not use the translocation event calls between different chromosomes. In order to understand, the effects of structural variations (SVs), we first grouped the deletions, inversions or duplications based on the length of the SVs:

*“Short-range SVs”* are identified as events with length of less than 2Mb and we mainly focused on these events in this study. Structural variations that are inflicting the TAD boundaries (SVIB) are identified as SVs that span through the whole length of a TAD boundary, rest of the SVs are classified as ‘within TAD’ in Figure 1B. To determine the distribution of random SVIB event, we used same bootstrapping method mentioned above, mainly generated random boundary events 10,000 times and calculated random SVIB event distributions. Z-scores and p-values were calculated based on observed number and distribution obtained from bootstrapping. In this study, we analyzed each event separately for deletion, duplication and inversions calls, albeit in a given sample these events might occur concurrently.

*“Long-range SVs”* are identified as event with length of more than 2MB and we mentioned the results obtained with long-range SVs inside the main text, when applied. In order to understand the germline SVIB occurrences, we downloaded structural alteration calls from three different studies: Abyzov et al,^28^ called deletion events (total of 8941) from 1000 genome project whole-genome sequencing data; Sudmant et al,^29^ called deletions (total of 7511) and duplications (total of 7501) from whole-genome sequencing data from 236 individuals representing 125 various human populations; Zarrei et al,^27^ performed a comprehensive review of deletions (total of 11530) and duplications (total of 1170) events in 23 different studies from 2647 different individuals. We noticed that number of SVIBs present in germline deletions and duplications are rare and happen less than expected by chance which was estimated with bootstrapping method.

We next profiled short-range SV and SVIB for each cancer studies in our ICGC dataset. To calculate average SV or SVIB per sample for each cancer studies, we divided the sum of all observed short-range SVs or SVIBs in a given cancer type to the total number samples in that cancer study. Observed SVs and SVIBs across cancer studies were plotted as stacked bar charts representing deletions, inversions and duplications with different colors.

To identify the recurrently affected boundaries in each cancer study, we generated a matrix where each column represents a sample in the cancer study and rows represent TAD boundaries. A binary score was assigned to each row (a TAD boundary) whether that boundary is affected by SVIB(s) in a given sample, results plotted in Figure 2c-d. Boundaries that are affected more than 10% of the samples in a cancer study, reported as recurrently affected boundaries in Supplementary Table 2. Median length of SVs per cancer type was calculated for all observed short-range SVs in each cancer types and plotted with the standard deviation of lengths in Supplementary Fig 3C. Constitutive insulated neighborhoods were obtained from Supplementary Table 8 of Hnisz et al.^51^ and SVs affecting only one anchor (CTCF-binding site) of an insulated neighborhood were considered as loop-disrupting SVs.

We determined flanking domain annotations of SVIBs, by identifying the type of the nearest domain for each SVIB break-ends. This analysis resulted in a half-matrix containing observed frequencies of pair-wise flanking domain types. We plotted the observed values for SVIBs deletions, inversion or duplications separately. To understand the genomic distribution of domain neighborhoods, we counted the flanking domains of each TAD boundary and plotted the obtained matrix in Supplemental Figure 3g.

We identified the nearest genes to the break-ends of SVIBs, as the nearest RefSeq genes that are not overlapping with the break-ends. RefSeq gene table was downloaded from UCSC genome browser on May 2016. We called the gene located up-stream of the 5’ end of an SV as “up-stream” gene and gene located down-stream of the 3’ end of an SV as “down-stream” gene for each SVIB. Expression fold-changes for each “up-stream” and “down-stream” genes were calculated by dividing observed normalized RPKM values in the particular sample with SVIB, with average normalized RPKM values in the rest of the same cancer study samples. We filtered the genes with low expression values (<0.1 FPKM), as fold changes with those genes would be seemingly high for even small fluctuations. Copy number variations could be another confounding factor for observed gene expression fold-changes. Therefore, we obtained consensus copy-number calls for the ICGC cohort based on consensus SV results. We removed the cases where copy numbers are more than four for either the “up-stream” and “down-stream” genes.

We used pyvcf (<u>https://pyvcf.readthedocs.org</u>) for loading .vcf files and pybedtools^54^ to perform genomic interval operations.

### Mutation Distributions

We downloaded the ICGC consensus calls for single nucleotide variations. In our analysis, we included ‘gray listed’ samples with the annotation of ‘false positive’ or ‘rescued’ in the comments column. We binned the mutations in 25 Kb non-overlapping windows along the genome, excluding Y and mitochondrial chromosomes. Mutation profiles were generated for each individual sample, sum of mutations in every cancer studies and sum of all mutations across the whole ICGC cohort. Next, we calculated the total number of mutations and sum of replication timing within each TAD, and normalized the obtained sums with the given TAD’s length.

For aggregate plots, mutation profiles and average replication timing in 25 Kb-windows from 500Kb upstream to 500Kb downstream of each TAD boundary demarcating low-active to low domains; low to low domains or low-active to low-active domains were plotted in Figure 4c. Similarly, average values of mutation load and replication timing for low to low-active domain were plotted in Supplementary Fig 4c, for repressed to active domains in Supplementary Fig 4d and repressed to low-active domains in Supplementary Fig 4e.

MSI status of each sample was inferred from the overall mutation load by taking the sum of mutations in 300Kb windows (between upstream 400 to 100 Kb) and (between downstream 100 to 400 Kb) for the TAD boundaries delineating low-active to low domains. We then used a linear regression model to identify samples that have no difference between upstream and downstream 300Kb mutation loads. We used a negative constant (-0.5 for mutations in 300 Kb window) as a filter to remove samples with low mutation loads such as leukemia or lower grade glioma samples. Colon cancer (COAD-US) MSI classifications were obtained from a previous study^42^ (Extended Data Figure 4j). Shinbrot et al^49^ identified Polε deficient cancers, (Supplementary Table 1a) by listing samples with Polε ultra-mutations (Polε-mutated MSS samples) and Polε mutated MSI samples. Previous studies^42,55^ suggested that, Polε mutated MSI samples have similar mutational load with MSI but Polε wild-type samples. Here, we noted that not only the number of mutations but also distributions of mutations around the TAD boundaries are comparable between Polε mutated MSI and MSI samples with wild-type Polε. In contrary, Polε mutated MSS samples contain higher mutation levels but distributions of those mutations around TAD boundaries are similar to the other MSS samples. Aggregate mutation loads around the TAD boundaries for MSI but Polε wild-type, MSI and Polε mutated, Polε mutated MSS samples were plotted in Supplementary Fig 4f.

### Mutation calling for inherited bi-allelic mismatch repair deficient (bMMRD) samples

Whole genome sequencing data of primary, recurrent brain cancers and normal control from a patient with bMMRD syndrome are available from Shlien et al. study^48^. Alignment files (bam) were downloaded from EGAD00001000369 and we used Mutect version 1.1.7^56^ for identifying mutations in primary (PD8640a) and recurrent (PD8640c) samples by using the default parameters. Later, identified mutations were filtered if the mutation is supported by at least 4 reads (t_alt_count > 4) in tumor sample and no reads in control (n_alt_count = 0). This filtering led to ∼716 thousand mutations in the primary sample and ∼267 thousand mutations in the recurrent sample. Mutation load around the TAD boundaries demarcating low to low-active boundaries were plotted for both samples in Supplementary Fig 4f.

## Acknowledgements

We thank the patients and their families for contributing to this study. We thank Sharon Dent, Zeynep Coban Akdemir, Tony Gutschner, Denise Spring, Jan Korbel and Josh Stuart for their critical readings of this manuscript. Special thanks to Adam Shlien for the bi-allelic mismatch repair deficiency sample access. We also would like to thank the discussion of Samir Amin, Sahil Seth, Floris Bartel, Tommy Mang, John Zhang. Authors are grateful to all ICGC subgroup participants for generating readily accessible mutation calls and uniformly analyzed gene expression dataset. This work was supported by a Cancer Prevention Research Institute of Texas award (R1205) and the Welch Foundation’s Robert A. Welch Distinguished Chair Award (G-0040 to P.A.F).

